# Cyclic-polymer grafted colloids in spherical confinement: insights for interphase chromosome organization

**DOI:** 10.1101/2023.04.05.535721

**Authors:** Jarosław Paturej, Aykut Erbaş

## Abstract

Interphase chromosome structures are known to remain segregated in the micron-sized eukaryotic cell nucleus and occupy a certain fraction of nuclear volume, often without mixing. Using extensive coarse-grained simulations, we model such chromosome structures as colloidal particles whose surfaces are grafted by cyclic polymers. This model system is known as Rosetta. The cyclic polymers, with varying polymerization degrees, mimic the functionality of structural protein complexes, while the rigid core models the chromocenter sections of chromosomes. Our simulations show that the colloidal chromosome model provides a well-segregated particle distribution without specific attraction between the chain monomers. Notably, linear-polymer grafted particles also provide the same segregation scheme. However, unlike linear chains, cyclic chains result in less contact between the polymer layers of neighboring chromosome particles, demonstrating the effect of DNA breaks in altering genome-wide contacts. As the polymerization degree of the chains decreases while maintaining the total chromosomal length (the total polymer length per particle), particles form quasi-crystalline order, reminiscent of a glassy state. This order weakens for polymer chains with a characteristic size on the order of the confinement radius. Our simulations demonstrate that polymer systems can help decipher 3D chromosomal architectures along with fractal globular and loop-extrusion models.

## Introduction

Cyclic or loop polymers are distinct from their linear counterparts due to their lack of free ends. Unlike linear chains, cyclic polymers are more compact and less prone to mix than their linear counterparts. Due to these conformational and topological differences, cyclic polymers exhibit lack of a relaxation plateaus in the viscoelastic behavior,^1,2^ lower interpenetration and frictional forces if coated on surfaces,^3–7^ and absence of topological entanglements.^8,9^ These unique properties of cyclic polymers enable them as polymeric components in stateof-the-art polymer-base materials^10^ while directing more attention towards the biological systems with naturally occurring cyclic-polymer structures.^11^

In materials science, colloids grafted by linear polymers (i.e., polymer brushes) are common practice to avoid colloidal accumulation by harvesting the limited interpenetration between polymer brushes.^12,13^ If such colloids are confined in spherical geometry, the packing conditions force them to organize according to the dimension of the confinement^14–16^ and allow them to exhibit properties that are not conceivable in bulk systems. Recent computational and experimental studies demonstrated that cyclic-polymer grafted surfaces are more effective in separating two polymer-grafted surfaces,^3,4^ offering new ways of tuning such colloidal systems, mainly by changing the amount of surface-grafted polymers or the polymerization degree of grafted chains.

From a biological perspective, cyclic polymers play an important role in understanding the mesoscale organization of genome.^9,17^ The genome is partitioned into chromosome structures, which are supramolecular protein-DNA complexes. In eukaryotic cells (e.g., mammalian cells), multiple chromosomes are isolated from cellular cytoplasm by a cell nucleus. Inside the micron-size nucleus, each chromosome occupies a certain volumetric region which is called chromosome territory.^18,19^ The observation of chromosome territories in a wide variety of species is explained by topological properties of cyclic homo-polymers. ^17,20^ However, chromosomes are of highly heterogeneous structures due to the cumulative effect of a dense array of structural proteins and nucleotide sequence; Often chromosomes form gene-poor, solid-like core regions,^21^ referred to as chromocenters. The spatial distribution of chromocenters inside the cell nucleus seems to be more regular than random for a wide range of species.^22,23^ In addition, the 3D organization of the chromosomes is further facilitated by the formation of loops of various sizes through the action of protein machinery, including structural maintenance of chromosomes (SMC) proteins.^24–26^ Given that the average loop size in chromosome structures is around several mega-base pairs,^27,28^ a single chromosome with close to 100 million base pairs can bear ∼100 loop structures. On a very coarse-grained level, each chromosome can be viewed as a complex of a large number of cyclic polymers emanating from a compact chromosome core (i.e., chromocenter). This depiction rather resembles a colloidal particle grafted with cyclic polymers. Therefore, the micron-scale segregation properties of chromosomes within the cell nucleus could share physical mechanisms similar to that of colloidal systems at a first-order level. Notably, such structures are also known as rosette polymers,^29,30^ and according to previous computational models for a fixed polymer size, ^31–33^ they can contribute to the segregation and organization of chromosomes. More recent studies have revealed that cellular concentration of loop-forming SMC proteins affects chromosome organization and even generates a distinction between various species.^26,34,35^ This suggests that the degree of polymerization or/and the effective grafting density of cyclic polymers, both of which can be controlled by structural nuclear proteins, can provide a polymer-physics perspective on how the genome is arranged within a nearly spherical nucleus that is roughly the size of several tens of microns.

The hypothesis presented in this study is that the presence of cyclic polymers on spherical particles causes differences in the way the particles are organized in confined spaces, compared to their linear counterparts. This is attributed to variations in their conformation and topology. By considering various grafting densities and polymerization degrees of grafts, we analyze the organization of at least n = 10 such particles at melt concentration by using coarse-grained molecular dynamics (MD) simulations. Our simulations indicate that even though linear and cyclic polymer grafted particles are organized similarly in 3D within a rigid spherical volume, cyclic polymers exhibit concentration profiles that are more uniform, and the grafted chains are less mixed. While these results are in line with the previous studies for chromosome organization at polymeric volume fractions of 10%,^31–33^ our comparison with linear chains at higher volume fractions also underlines the effect of chain topology at biologically-relevant volume fractions.^36^

## Simulation model

Using a coarse-grained model we performed molecular dynamics simulations of polymer-grafted colloidal particles in spherical confinement. To simulate polymers, the bead-spring model of Kremer and Grest was utilized.^37,38^ A polymer chain was composed of N spherical beads of equal size and mass connected by massless springs. Both bonded and non-bonded monomer-monomer pairs repel each other by the truncated and shifted Lennard-Jones (LJ) potential

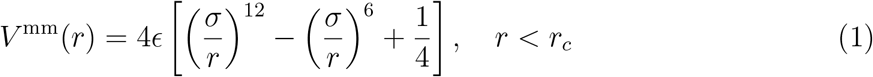

with the cutoff radius *rc*=2^1/6^*σ* and *V* ^mm^(*r* ≥ *r*_*c*_) = 0. In Eq. (1) *ε* is the strength of the LJ interaction and σ is the monomer diameter which is taken as the units of energy and length, respectively. Correspondingly, the units of volume fraction, temperature and time are [*ϕ*] = *σ*^−3^, [*T* ] = *ϵ*/*k*_*B*_ and [*τ*] = (*mσ*^2^/*ϵ*)^1/2^, where k_B_ is the Boltzmann constant and m is the monomer mass. The bonds between subsequent beads along the chain are mimicked by non-linear springs

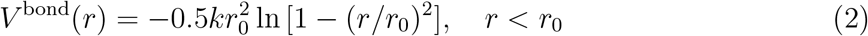

with *V* ^bond^(*r* ≥ *r*_*0*_) = *∞, r*_0_ = 1.5*σ* and *k* = 30*ϵ*/*σ*^2^. A polymer-grafted colloid was represented as a spherical core with f attached chains. The diameter of the colloidal particle is *σ*_C_ = 5*σ* and its mass 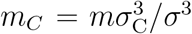. In order to account for interactions between the polymer beads with colloids, which are incompatible in size, we incorporate the so-called expanded LJ potential^39^

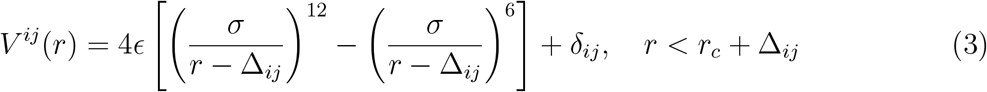

with *V* ^*ij*^(*r* > *r*_*c*_ + Δ_ij_) = 0. In the above equation the quantity *δ*_ij_ is such that *V*^*ij*^(*r*_*c*_ + Δ_ij_) = 0. The potential shift Δ_ij_ for monomer-colloid interaction is Δ_mC_ = (*σ*_C_ *- σ*)/2 whereas for the colloid-colloid interaction is Δ_CC_ = *σ*_C_ − *σ*. Finally, the spherical confinement with radius R was modeled as a structureless, impenetrable, and repulsive rigid wall. We assume the monomer-wall interaction of the same form as in Eq. (1). The only difference is the replacement of the inter-particle distance r by 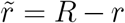 in spherical coordinates, where the origin of the system coincides with the center of the sphere.

The simulations were conducted for n = 10 confined colloids and linear or cyclic polymer chains were attached to them with different arrangements as shown in Fig. 1a. While keeping the total number M of monomers in the systems constant, we performed simulations with different values of the degree of polymerization N of individual chains ranging from 15 to 240, as well as varying the number f of attached chains from 10 to 160. Initially, polymer-grafted colloids were distributed randomly inside a spherical volume with a large radius *R*_*i*_ ≈ 127*σ* corresponding to a density (i.e., volume fraction) 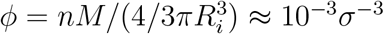. Subsequently, the radius was gradually decreased to *R* ≈ 27*σ* to achieve the target density *ϕ* ≈ 0.3*σ*^−3^ (i.e., 30% polymer content), cf. Fig. 1b. Finally, simulations were followed by a production run lasting 10^6^*τ*.

**Figure 1:**
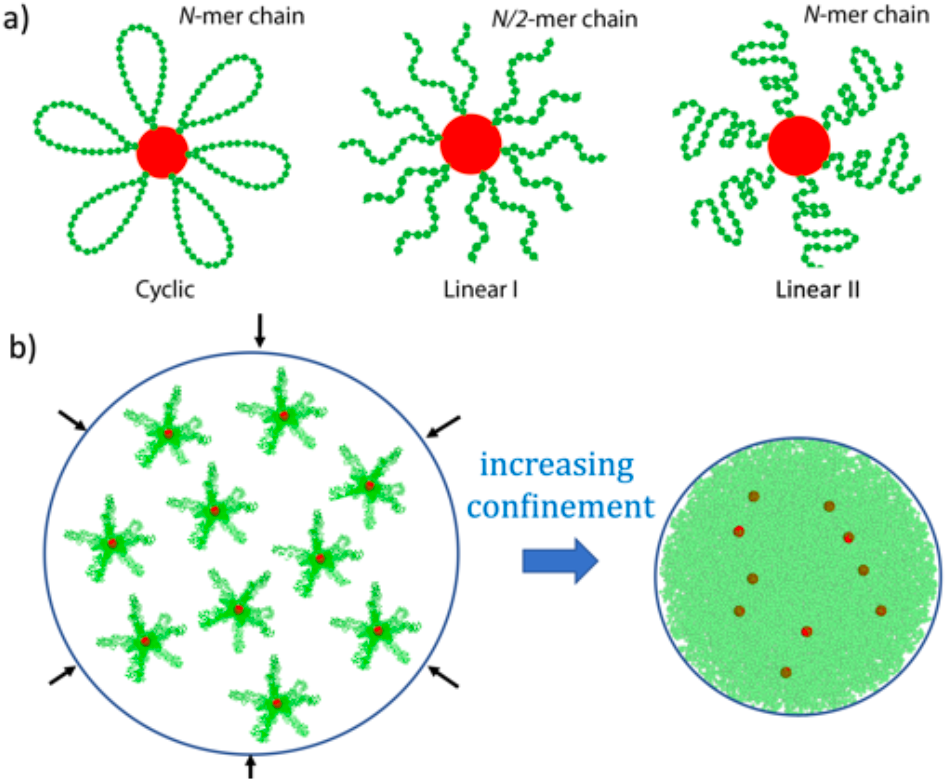
a) The molecular structure of the chromosome models consists of polymers that are grafted onto a colloidal particle. The model chromosomes are packed inside a sphere with an initial radius, *R*_*i*_ = 127*σ*. The volume of the sphere is decreased to *R* = 27*σ* to obtain the prescribed polymer volume fraction of *ϕ* = 0.3*σ*^−3^. Each spherical volume contains *n* = 10 colloidal particles. The total number of chain monomers per colloidal particle is always fixed at *M* = 2400 but the distribution of these monomers among the grafted chains varies based on the number *N* of monomers per chain in each case.

The simulations were conducted using the Large-scale Atomic/Molecular Massively Parallel Simulator (LAMMPS).^40^ The velocity Verlet algorithm was used to integrate Newton’s equations of motion, employing a time step Δ*t* = 0.005*τ*. The temperature T was maintained by the Langevin thermostat with a friction coefficient *ζ* = 0.5 m*τ*^−1^. The simulation snapshots were rendered using the Visual Molecular Dynamics (VMD).^41^

A direct one-to-one mapping of our models to chromosome organization is not possible due to the simplicity of the coarse-grained model. However, if each bead is assumed to represent roughly 10^4^ base pairs, *M* = 2400 corresponds to 2.4 × 10^6^ base pairs, and for *n* = 10 particles, our model represents more than 10^7^ base pairs. While this number is small compared to human chromosomes with ca. 100 million base pairs, it is statistically large enough to obtain Gaussian statistics for our chains.^8,37^ Further, this polymer size provides an optimum interphase-chromosome volume fraction of around *ϕ* 30%, which is consistent with the recent electron microscopy studies^36^

## Results

In our simulations, we model chromosomes as colloidal particles, with each particle having f cyclic (or linear) flexible chains grafted onto it, where each chain is composed of *N* monomers (cf. Fig. 1). To create different structural configurations, we vary *f* and *N* individually while keeping fixed the total number *M* = *fN* of chain monomers per particle fixed and the overall particle concentration *ϕ*. In this way, we can simulate different partitioning scenarios of a single chromosome into varying numbers of loops, as observed in the cell nucleus,^26^ which is controlled by topological proteins such as SMCs. Our analysis considers the organization of *n* = 10 colloidal chromosomes inside a rigid, non-penetrable spherical shell with a radius *R ≈* 27*σ* mimicking either a cell nucleus or colloidal confinement. This confinement results in a volume fraction of *ϕ* = 0.3*σ*^−3^.^36^ The functionality *f* is related to the grafting density according to the equation:

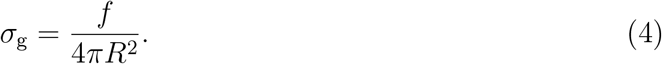

Our objective is to identify the optimal conditions, specifically the appropriate values of *σ*_g_ and *N*, that would promote the segregation of colloidal particles coated with cyclic polymers throughout the entire cell nucleus.

In Fig. 2, we display the organization of polymer-grafted particles with different *f* and *N* values for various chain topologies. The snapshots depict the colloidal (core) particle (represented by red spheres) onto which polymers are attached. These cores can model either the chromocenters, which exhibit properties of a polymeric solid^21^ or metal nanoparticles.^16^ We observe the segregation of colloidal particles grafted by cyclic polymers such that the core particles have no steric contact with each other. This means that the colloidal particles are kept separate from one another by the outer layers (shells), which are composed of polymers grafted onto their surfaces. However, the extent of overlap between the polymer shells varies depending on the value of *f*. Through visual examination, we observe that for systems with longer rings (i.e., smaller *f*), there is a higher degree of interpenetration between the outer layers of adjacent colloids. In fact, this trend depends on the grafting density and polymerization degree. For high grafting densities and low polymerization (e.g., *f* = 80 and *N* = 30), polymer shells overlap, but individual polymer-grafted particles are of hairy particle morphology in appearance. Consequently, mixing between polymer shells is visually absent (Fig. 2a). On the contrary, for low grafting densities and large polymer lengths (e.g., *f* = 10 and *N* = 240), the polymer shells of neighboring colloids can overlap significantly (cf. Fig. 2a). The pervaded volume of polymers fills the spherical volume while the polymer of one particle can interpenetrate through the shells of neighboring particles. Notably, as *f* decreases (and *N* increases), the structures of individual polymer-coated particles tend to become more spherical since shorter and densely grafted chains are more likely to adopt a stretched conformation.^42^ The presence of shorter chains in these hairy particles can result in a glassy behavior, which makes them less susceptible to diffusion on average.^**?**^ It is noteworthy that the stiff chromosome structures also change the collective morphology of all confined chromosomes. Specifically, the spherical shape changes from being perfectly round to having protrusions, and empty spaces emerge between the chromosomes and the boundary of the spherical confinement.

**Figure 2:**
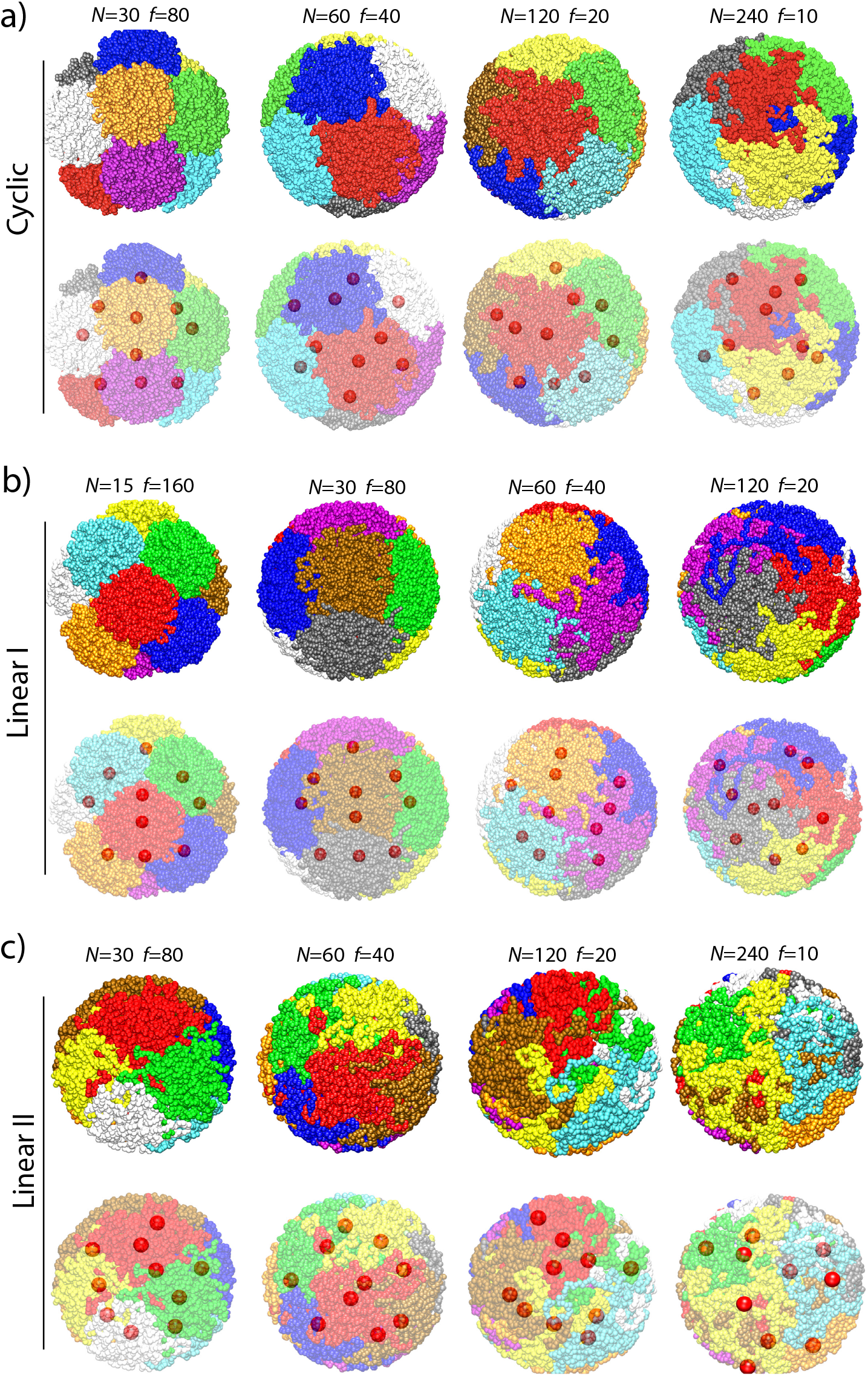
Representative molecular dynamics snapshots of polymer-grafted colloid organized in spherical confinement. The chain monomers of each polymer-grafted particle are color-coded for clarity. The bottom frames in each case display the chains in a transparent mode to reveal the core particles, depicted as red spheres. The parameters *N* and *f* correspond to the polymerization degree of grafted chains and the number of grafted chains per particle, respectively. Each spherical volume contains *n* = 10 colloidal particles.

To observe the effects of cyclic topology, we also investigate systems of colloidal particles grafted by linear polymers. To construct the linear systems while keeping the total number of beads per core identical, we follow two strategies: i) we cut the *N*/2th bond of a cyclic polymer and effectively double the functionality while halving the chain length per core (Fig. 1a), ii) we replace each cyclic chain by a linear chain without changing its functionality (Fig. 1a). We refer to these two models as Linear I and Linear II. In these simulations, the arrangement of colloids exhibits comparable patterns to those observed in the cyclic-polymer systems in relation to *f* (or *N*) (Fig. 2b,c). However, a visual inspection reveals that polymer layers in those two cases can mix with each other more drastically The most drastic change occurs for the Linear II cases, in which even the highly functionalized colloids do not display a quasi-lattice order that is observed for Cyclic and Linear I cases (e.g., *N* = 30-case shown in (Fig. 2c).

Next, to quantify the amount of overlap between colloids, we calculate the number of steric contacts between polymer chains of neighboring colloids by assigning a contact distance of *r*_*bb*_ = 1.5*σ* between the beads of any two colloidal particles (Fig. 3). This distance corresponds to the occurrence of a steric contact between any two beads (cf. Eq. 1). This calculation confirms the effect of increasing contacts with increasing *f* (and decreasing *N*) (Fig. 3). However, the number of contacts between the beads of cyclic chains is systematically lower than those of linear cases (Fig. 3). Since both linear and cyclic cases have an equal number of beads per colloid, and *f* and *N* are related via *M* = *N*/*σ*_*g*_ for a fixed *M*, the difference between the linear and cyclic cases in Fig. 3 indicates that the main effect controlling the number of contacts is the chain topology. This significant difference is due to the lack of free ends in the cyclic polymers that are known to suppress the inter-digitation between two polymer brushes composed of cyclic polymers.^3^ Notably, in Fig. 3 we observe a saturation behavior for large *N* regardless of the topology. The trend remains unchanged even when plotted on a logarithmic scale (shown in the inset of Fig. 3). We attribute this behavior to the finite size of the confinement, which is *R* = 50*σ*; as the size of the polymer chains increases, the walls of the confinement can restrict the amount of inter-polymer contacts.

**Figure 3:**
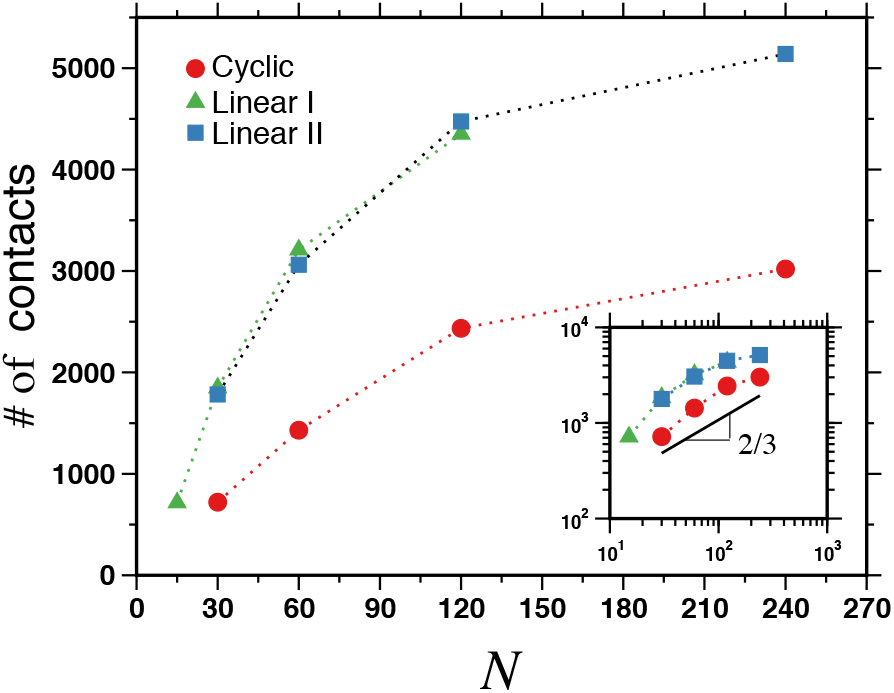
The average number of pairwise contacts between the shell beads of adjacent colloids for linear and cyclic grafts defined in Fig. 1. The inset shows the same data on a logarithmic scale.

The number of contacts between the chains of two colloids could be related to the polymerization degree of grafted chains by utilizing the scaling arguments originally suggested for two weakly interacting planar brushes.^43^ The thickness of the overlap region between the polymer shells of two neighboring colloids separated by a distance *d* can be expressed as 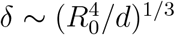 where *R*_*0*_ ∼ *N*^*ν*^ is the characteristic size of the polymer chain, and 1/*ν* is the corresponding fractal dimension of the polymer chains. Assuming that the inter-particle distance *d* ∼ *ϕ*^1/3^ is constant, we can write

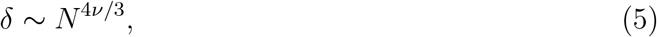

where *ν* = 1/2 for linear polymers and *ν* = 1/3 loop polymers,^2,20^ respectively. If the number of contacts is assumed to increase with the overlap distance, both linear and cyclic polymers exhibit an increase as *δ ∼ N* ^2/3^ vs. *δ ∼ N* ^1^, respectively. We should note that for cyclic polymers *ν* = 1/3 exponents usually appear for longs chains (e.g., *N* > 1000)^2,17,44^ and therefore, for our chain sizes, *ν* = 1/2 could be a more valid approximation. Nevertheless, Fig. 3 demonstrates that the main suppressive effect leading to a lower number of steric contacts between the overlapping polymer shells of two colloidal particles is due to cyclic chain topology rather than the grafting density or polymerization degree.

Subsequently, we determine how the concentration profile of monomers within the spherical confinement is affected by the functionality of grafted chains and by their topology. In order to quantify the monomer distribution, in Fig 4 we plot the radial density profiles of monomers as a function of radial distance from the center of spherical confinement. For cyclic grafts, with the low interpenetration between the chromosomes with high grafting density and low polymerization (e.g., *f* = 80 chains and *N* = 30 beads per chain), the concentration at the center of the spherical volume is negligible. This low density implies the presence of polymer-free voids resulting from the tight arrangement of polymer-grafted particles. It is worth mentioning that similar voids have been observed in DNA-coated nanoparticles, which result in metallic-like particle phases.^45^ As *f* decreases (*N* increases), the monomers fill the voids and the concentration becomes uniform throughout the spherical volume. If we compare cyclic and linear topology, cyclic polymers tend to produce a relatively more non-uniform density profile, particularly near the spherical boundary (Fig 4). As the value of *N* increases, a slight bump in the concentration profile of cyclic polymers persists. This feature is either weaker or absent in the case of linear polymers. These findings support the notion of a higher density of colloidal particles near the boundary, as reported in earlier studies but at lower volume fractions.^31,32^

**Figure 4:**
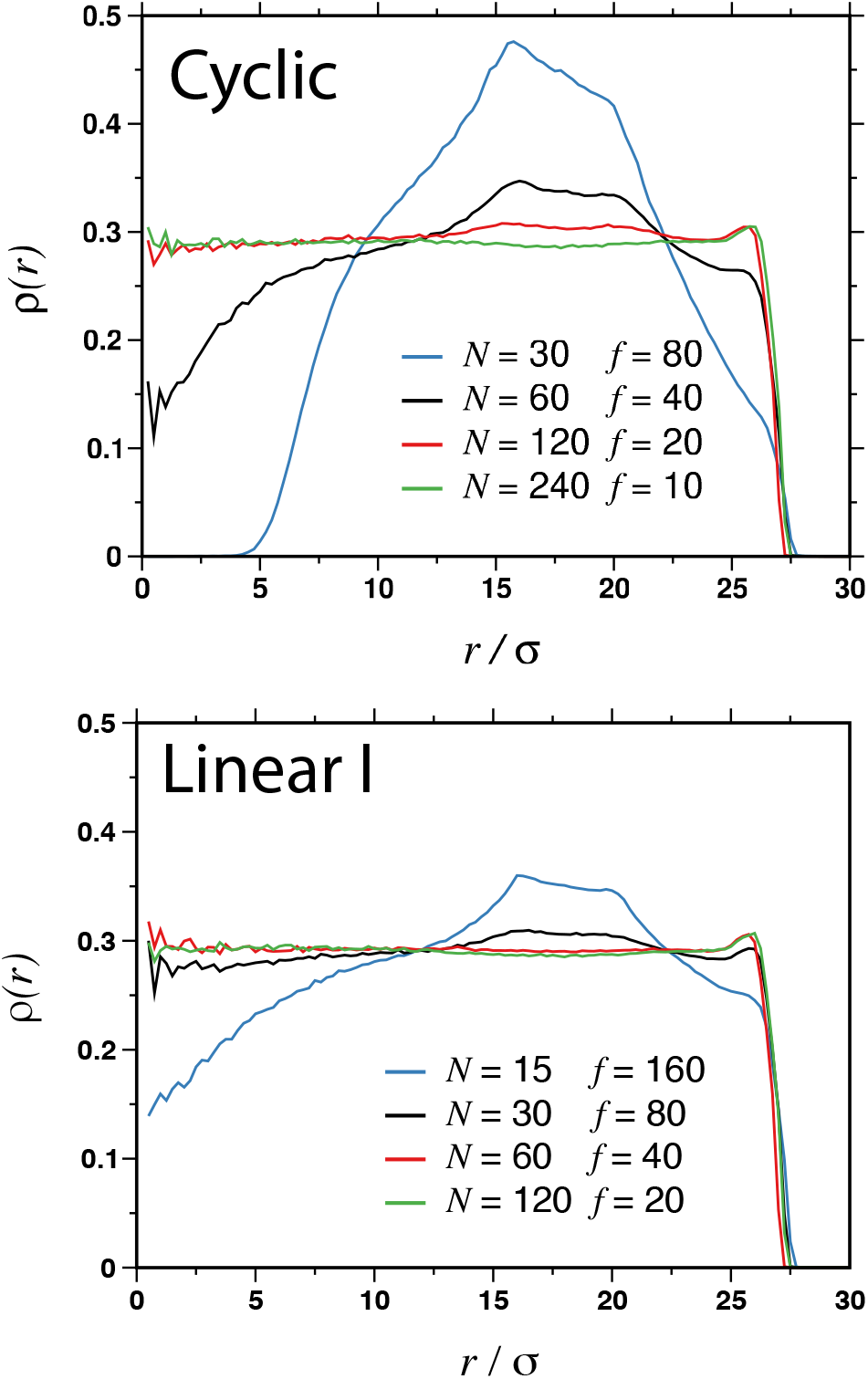
The radial concentration profiles *ρ*(*r*) of chain monomers as a function of the distance from the center of the spherical confinement with radius *R ≈* 27*σ*. The data displayed for two graft architectures with different functionalities *f* and degree of polymerization *N* as indicated.

In Fig. 2, polymer-grafted particles form any quasi-crystalline structure inside the spherical confinement for cyclic and linear-I cases. This order weakens as the functionality decreases (i.e., as *N* increases). In a cell nucleus, each chromosome has a different length (different number of DNA base pairs), thus, the emergence of such order may not be relevant as opposed to synthetic colloidal systems.^14^ Nevertheless, spermatozoan cells with highly-compact chromatin structures, which can correspond to high functionality and low polymerization in our model, exhibit discrete centromere positioning inside the nucleus.^23^ To quantify the relationship between such quasi-crystalline ordering and chain size, we calculate the average distance between the cores, *d*, and its distribution functions for various *N* and *f* cases for both cyclic and linear-polymer grafted particles (Fig. 5). While the inter-core distance often shows an average value *d* ≈ 26*σ* independent of grafting parameters and topology, the distribution of *d, ρ*_*core*_(*d*) depends on both topology (cyclic versus linear) and grafting parameters. For densely grafted chains (high *f*), the distribution has two humps, separated by a zero-concentration plateau, suggesting a BCC-like lattice structure. This data indicate low positional fluctuations for the colloids densely grafted by short chains,^46,47^ trapped by an effective cage imposed by identical neighboring particles. Additionally, each hump has two sub-peaks for the two shortest chain lengths (i.g., *N* = 15, 30 and *N* = 30, 60) separated by a distance *≈* 3*σ*. If f decreases (and N increases), the strength of the ordered structures observed in the *ρ*_*core*_(*d*) profiles decreases as well. This implies that the neighboring particles have a weaker caging effect.^47^ It is noteworthy that for our linear II cases, for which particles have the same functionality as in the cyclic case, the peaks are broader and lower, indicating also a weaker caging effect from linear chains in constraining particle positions.

**Figure 5:**
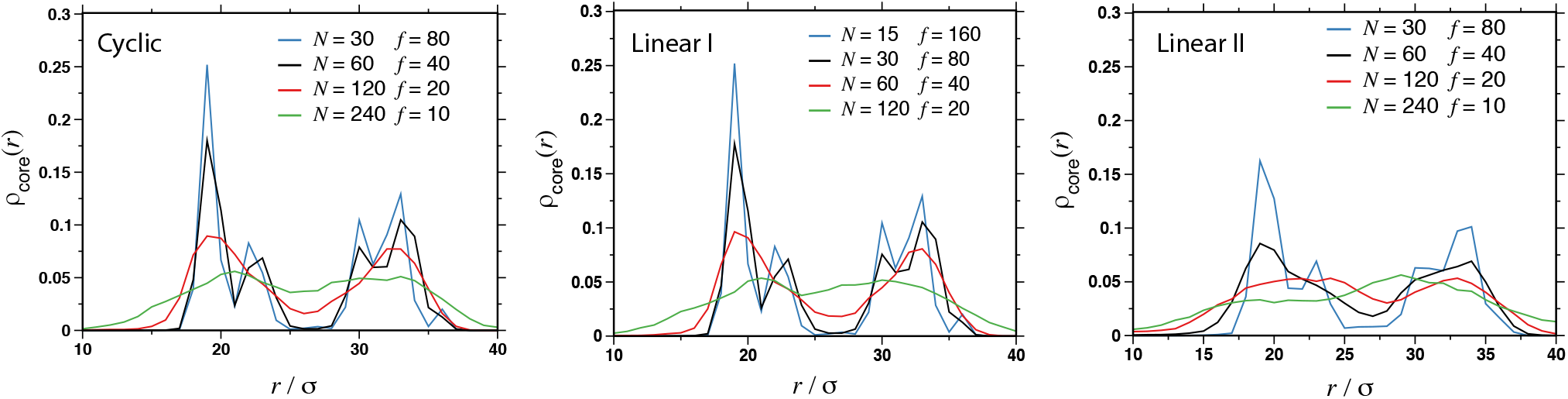
The normalized distributions *ρ*_*core*_(*r*) of chromosome positions within spherical confinement measured as a function of the radial distance *r* from the center of a sphere. Data displayed for different architectures of grafts defined in Fig. 1 and various functionalities *f* and degree of polymerization *N* as indicated in the legend.

## Discussion

In this work, we use coarse-grained polymer simulations to unveil polymeric contributions to the organization of polymer grafted colloidal particles in rigid spherical confinement. Particles grafted by cyclic and linear polymer chains show that both the polymerization degree and grafting density of the chains affect the organization. Simulations show that cyclic polymers can enhance the segregation (e.g., fewer contacts between chains) more strongly as compared to linear chains. While both of our short (i.e., *N* = 30 monomers) and long (i.e., *N* = 240) cyclic chains can cause this effect, long loops provide relatively more uniform polymer distribution in the confinement whereas shorter cyclic chains lead to a quasi-crystalline particle order inside the spherical confinement, similar to the behavior of similar colloidal particles in bulk.^47^ Our simulations with linear-chain grafted particles indicate that indeed this is not a simple polymer-size effect but rather a direct effect of the topology of cyclic polymers with no free ends. Nevertheless, the repulsion between the polymer layers of the particles depends on the grafting parameters and chain sizes in both cyclic and linear-polymer cases^3,43,48^

Our results could have important consequences to understand the nuclear organization of interphase chromosomes in micron-size eukaryotic cell nuclei. While chromosomes are highly complex supramolecular protein-DNA complexes, they can be considered hetero-polymers by embedding all protein-mediated effects in various pairwise interactions^49,50^ or polymer rigidity.^51^ In this view, some chemically distinct sections of a chromosome can be considered as in poor solvent conditions (e.g., chromocenters), thus collapsed, while other sections could be considered in their good-solvent conditions and thus relatively swollen. This depiction can allow us to consider each chromosome as a polymer-grafted micron-size colloidal particle, such that loops of various sizes emanate from a solid-like core (i.e., chromocenter).^21,25,27,31^ Hence, on the first-order level, micron-scale segregation properties of chromosomes inside the cell nucleus can share a similar physical mechanism with colloids in spherical confinement.

Remarkably, experiments and image analyses have shown that chromocenter distributions inside the cell nuclei of various mammals and plant species exhibit relatively ordered, almost regular organization patterns.^22,23^ In this context, our simulations suggest that entropic repulsion can contribute to such regular distribution of chromocenters while preserving chromosome territories.^31,32^ The lower number of contacts between the particles grafted by cyclic polymers can further contribute to this repulsion process, further ensuring the demixing of chromosome chains. This is in conjecture with the equilibrium globular model, in which chromosomes are modeled as loop polymers.^17,20^ However, equilibrium globular inevitably considers chromosomes as homopolymers and does not distinguish between the different conformational properties of chromosome sections (e.g., chromocenters). Further, chromocenters should preserve their segregation patterns after each cell division since this segregation is a criterium for the classification of species.^26,34^ The colloidal chromosome model presented here combines the sold-like nature of chromocenters and pericentromeric chromatin and the formation of loops by chromatin architectural proteins (i.e., SMC) and could contribute to the explanation of the recurrent segregation of chromocenters and chromosomes. ^22,23,26**?**^ Notably, previous studies rather demonstrate that chromocenter segregation is not provided by linear chains for volume fractions much less than *ϕ* 30%.^31^ We here show that chromosomes and their cores can stay separate even at *ϕ* 30%, which seems to be the optimum value for most cell types.^36^ At this concentration, linear-polymer grafted particles can also exhibit segregation properties, albeit with a higher number of contacts than cyclic proteins.

Our simulations also suggest that as the loop size increases (e.g., a lower activity or concentration of SMC or similar proteins), individual chromosomes can overlap and exhibit more positional fluctuations relative to each other. While shorter chains (i.e., *N* < 100) can segregate chromosomes well, chromosome structures can become effectively more *stiff*. Consequently, the overall shape of all the chromosome chains does not take the shape of the spherical confinement as expected from a polymer liquid. Such *stiff* chromosomes could generate protrusions and bumps on the surface of the elastic cell nucleus, within which they are confined. If the cellular levels of the proteins that are responsible for loop formation

Considering that SMC proteins are responsible for the formation of a dynamic loop topology,^25,28^ the expression levels of these proteins and their residence times can be correlated with the amount of loop per chromosome.^26,35^ In this case, it is conceivable to think of a scenario, in which the over-expression of these proteins (e.g., shorter loops) can change the shape of the cell nucleus. A future study with deformable shell models can enlighten the morphological effects of loop-forming proteins.

Lastly, our simulation with linear-polymer grafted colloids demonstrate that inter-chromosomes contacts could increase if chromosome loses their cyclic topology, and as more free ends form. DNA breaks tend to increase free DNA ends in disease and aging.^52^ Linear chains of a polymer layer tend to penetrate to the opposing layer stronger as compared to a layer of cyclic chains,^3^ as we also show here. As an overarching prediction of our simple coarse-grained simulations, we can speculate that such breaks can increase contact between the genes and favor the clustering of double-stranded DNA breaks^53,54^ that otherwise should not be in contact with each other in a healthy state.

## Conclusions

To summarize, our simulations demonstrate that polymerization degree and grafting density could be used to control the segregation properties of colloidal particles in confinements, and such systems can serve as simple model architectures to comprehend the complex nature of 3D genomic architecture in nuclear confinement. On one hand, the similar polymer-based model can serve to reveal large-scale organization patterns of the genome, on the other hand, they can increase the reach of polymer physics to problems beyond synthetic systems.

This work has been supported by the National Science Center, Poland (Grant Polonez Bis No. 2021/43/P/ST3/01833). We thank K. Haydukivska for their critical reading of the manuscript. We are also grateful to PL-Grid Infrastructure for a generous grant of computing time at the Prometheus cluster.

